# Assessing Spns2-dependent S1P Transport as a Prospective Therapeutic Target

**DOI:** 10.1101/2024.03.26.586765

**Authors:** Y Kharel, T Huang, K Dunnavant, D Foster, GMPR Souza, KE Nimchuk, AR Merchak, CM Pavelec, ZJ Juskiewicz, A Gaultier, SBG Abbott, J-B Shin, BE Isakson, W Xu, N Leitinger, WL Santos, KR Lynch

## Abstract

S1P (sphingosine 1-phosphate) receptor modulator (SRM) drugs interfere with lymphocyte trafficking by downregulating lymphocyte S1P receptors. While the immunosuppressive activity of SRM drugs has proved useful in treating autoimmune diseases such as multiple sclerosis, that drug class is beset by on-target liabilities such as initial dose bradycardia. The S1P that binds to cell surface lymphocyte S1P receptors is provided by S1P transporters. Mice born deficient in one of these, spinster homolog 2 (Spns2), are lymphocytopenic and have low lymph S1P concentrations. Such observations suggest that inhibition of Spns2-mediated S1P transport might provide another therapeutically beneficial method to modulate immune cell positioning. We report here results using a novel S1P transport blocker (STB), SLF80821178, to investigate the consequences of S1P transport inhibition in rodents. We found that SLF80821178 is efficacious in a multiple sclerosis model but – unlike the SRM fingolimod – neither decreases heart rate nor compromises lung endothelial barrier function. Notably, although Spns2 null mice have a sensorineural hearing defect, mice treated chronically with SLF80821178 have normal hearing acuity. STBs such as SLF80821178 evoke a dose-dependent decrease in peripheral blood lymphocyte counts, which affords a reliable pharmacodynamic marker of target engagement. However, the maximal reduction in circulating lymphocyte counts in response to SLF80821178 is substantially less than the response to SRMs such as fingolimod (50% vs. 90%) due to a lesser effect on T lymphocyte sub-populations by SLF80821178. Finally, in contrast to results obtained with Spns2 deficient mice, lymph S1P concentrations were not significantly changed in response to administration of STBs at doses that evoke maximal lymphopenia, which indicates that current understanding of the mechanism of action of S1P transport inhibitors is incomplete.

## Introduction

Study of the first S1P receptor modulator (SRM; fingolimod (FTY720)) revealed a theretofore unrecognized aspect of S1P biology, *i.e*., S1P is required for the correct distribution of immune cells; for example, egress of T lymphocytes from lymphoid tissues^1^. S1P1 receptor agonists such as the SRM drugs desensitize those receptors and thereby prevent their responding to extracellular S1P, which limits lymphocyte chemotaxis. This disruption of lymphocyte trafficking results in peripheral blood lymphopenia^1^, which is the pharmacodynamic index of SRM target engagement. While five SRMs are now approved medicines for treatment of multiple sclerosis and/or ulcerative colitis, this drug class has on-target liabilities such as 1^st^ dose bradycardia^2^ indicating a need to discover alternative interventions to manipulate S1P signaling.

The SRM mechanism of action suggests that reshaping extracellular S1P concentration gradients is an alternate strategy to interfere with S1P signaling for therapeutic benefit. S1P gradients are generated by the combined activities of exofacial lipid phosphatases (*e.g.*, Plpp3^3^), S1P transporters and catabolism of intracellular S1P by S1P lyase^4^ and S1P phosphatases^5^. Drug-like phosphatase inhibitors are notoriously difficult to find and S1P lyase inhibition is nephrotoxic^6^. While several S1P transporters have been proposed^7^, the best characterized to date are the solute carriers, Mfsd2b (SLC59A2) and Spns2 (SLC63A2). Forced expression of either protein results in S1P release by naïve cells^8,9^. Mfsd2b expression is reportedly limited to red blood cells (RBCs) and platelets^10^ while Spns2 is expressed by a variety of cell types including endothelial cells^11^, kidney pericytes^12^, activated monocytes^13^ and microglia^14^.

Spns2 null mice^15,16^ and endothelium-specific^17,18^ Spns2-deficient mice have significantly reduced (∼50%) peripheral blood lymphocyte counts compared to control mice, which is reminiscent of the response to SRM administration. Spns2 null mice also have negligible S1P in thoracic duct lymph compared to control mice^17,18,19^. Together, these facts suggest that STBs (S1P transport blockers, specifically, Spns2-dependent S1P transport) and SRMs have similar mechanisms of action.

We have discovered small molecule inhibitors of Spns2-dependent S1P release and use these to discern the extent to which STBs mimic the actions of SRMs. Using STBs on distinct chemical scaffolds^20,21^, we discovered that these compounds drive a dose-dependent decrease in blood lymphocyte counts in mice and rats^20,21^. Further, we found that our prototype STB, SLF1081851, is efficacious in models of renal fibrosis^12^. Herein, we report on SLF80821178, which is representative of a third STB structural class. We found that SLF80821178 is efficacious in a standard model of multiple sclerosis, and we used this molecule to investigate potential toxicities associated either SRMs or genetic manipulation of *Spns2* in mice. We also used SLF80821178 and our earlier STBs to investigate the mechanism of action of these inhibitors.

## Results

### Compound Characterization

SLF80821178 was obtained by replacing the oxadiazole-propylamine moiety of SLF1081851^20^, with a urea-piperazine group (refer to Fig. 1 for structures). This substitution resulted in increased oral bioavailability as well as a nearly 50-fold increase in potency over SLF1081851 in an assay of Spns2-dependent S1P release from HeLa cells (IC_50_ 1.9 µM^20^ vs. 0.05 µM). Like previously reported STBs^20,21^, single dose administration of SLF80821178 to mice resulted in a dose-dependent decrease in circulating lymphocytes with a maximal decrease of ∼50%. The details of the characterization of SLF80821178, a structure-activity relationship of compounds on the urea-based scaffold and their synthetic routes are the subject of a separate report (Foster, *et al*., submitted for publication).

**Figure 1:**
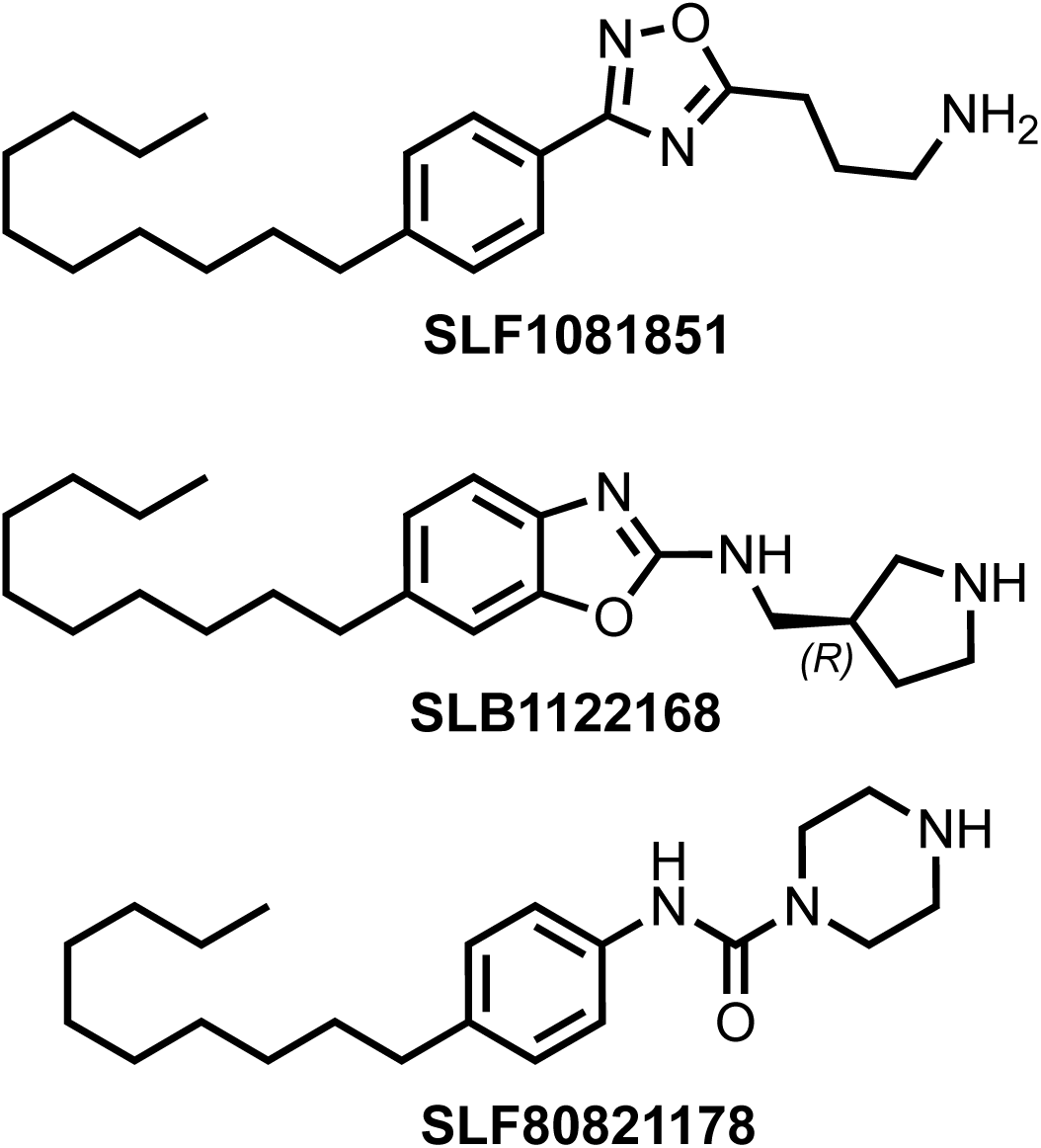
Structures of STBs used in this study. SLF1081851^17^ (3-(3-(4-decylphenyl)-1,2,4-oxadiazol-5-yl)propan-1-amine hydrochloride), SLB1122168^18^ ((*R*)-6-decyl-*N*-(pyrrolidin-3-ylmethyl)benzo[*d*]oxazol-2-amine hydrochloride), SLF80821178 [Foster, *et al*. submitted] (*N*-(4-decylphenyl)piperazine-1-carboxamide hydrochloride)

In investigating the selectivity of SLF80821178, we found no effect on either release of S1P from mouse red blood cells (RBCs) *ex vivo* at concentrations up to 5 µM (Fig. 2A) or on plasma S1P concentrations in SLF80821178 treated mice (see below). These results indicate that the compound is not interfering with S1P release by RBCs, which is Mfsd2b-dependent^10^. Further, SLF80821178 neither promoted nor blocked S1P-evoked migration of mouse spleen T-lymphocytes (Fig. 2B), which indicates a lack of engagement of the S1P1 receptor by SLF80821178 on those cells. Finally, SLF80821178 inhibited recombinant sphingosine kinase (SphK1, SphK2) activity only modestly at a concentration of 10 µM (Fig. 2C).

**Figure 2:**
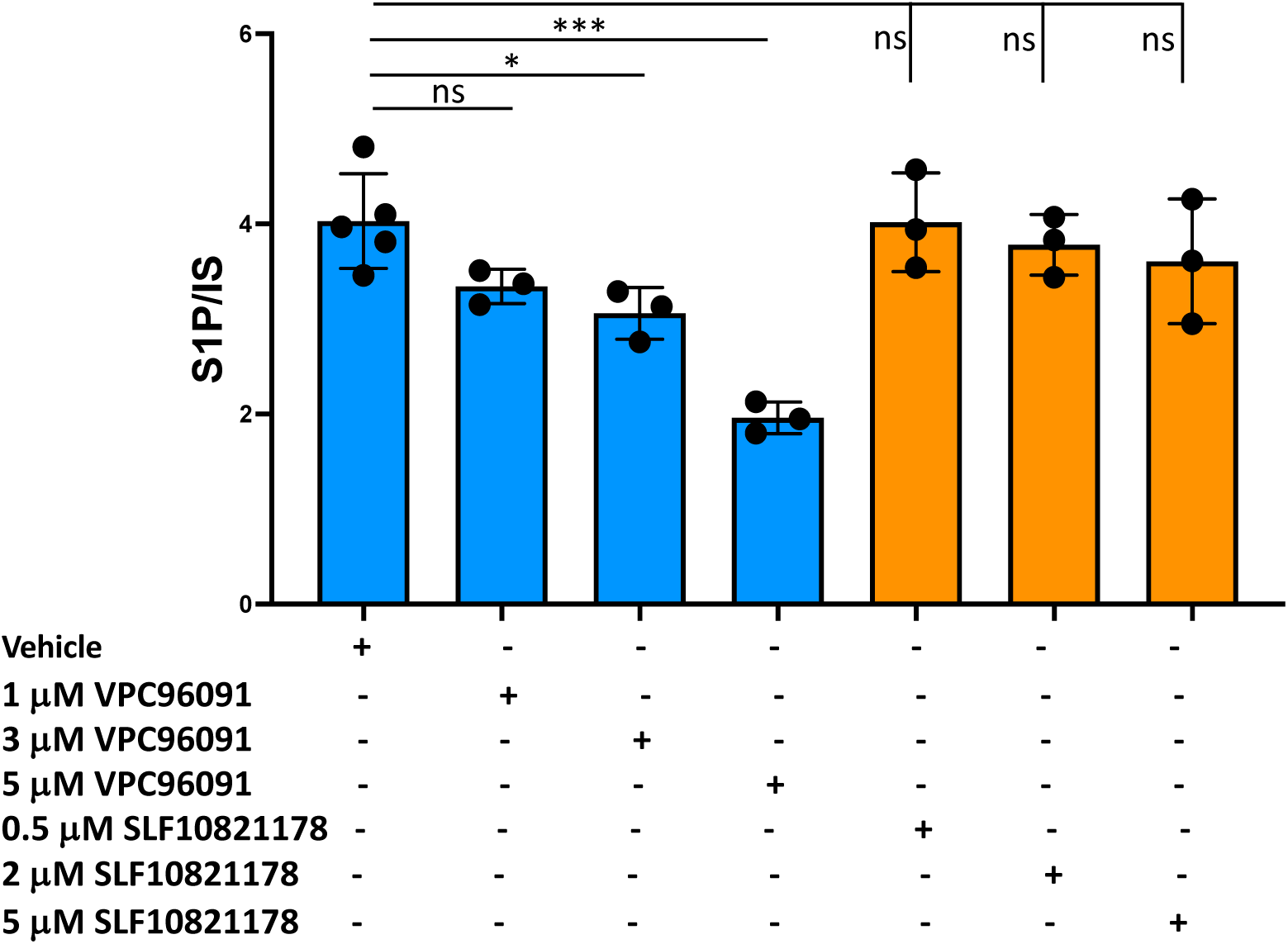
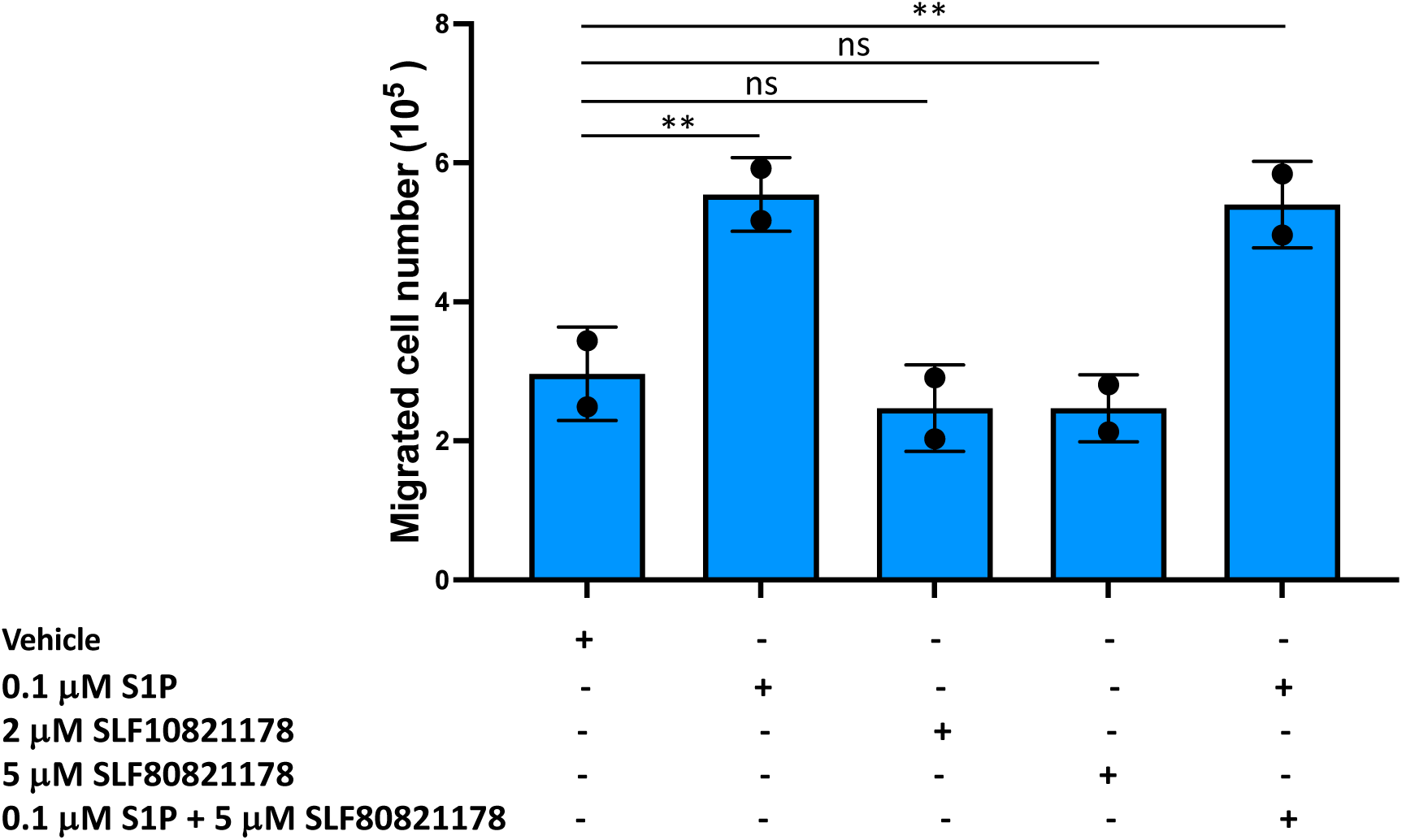
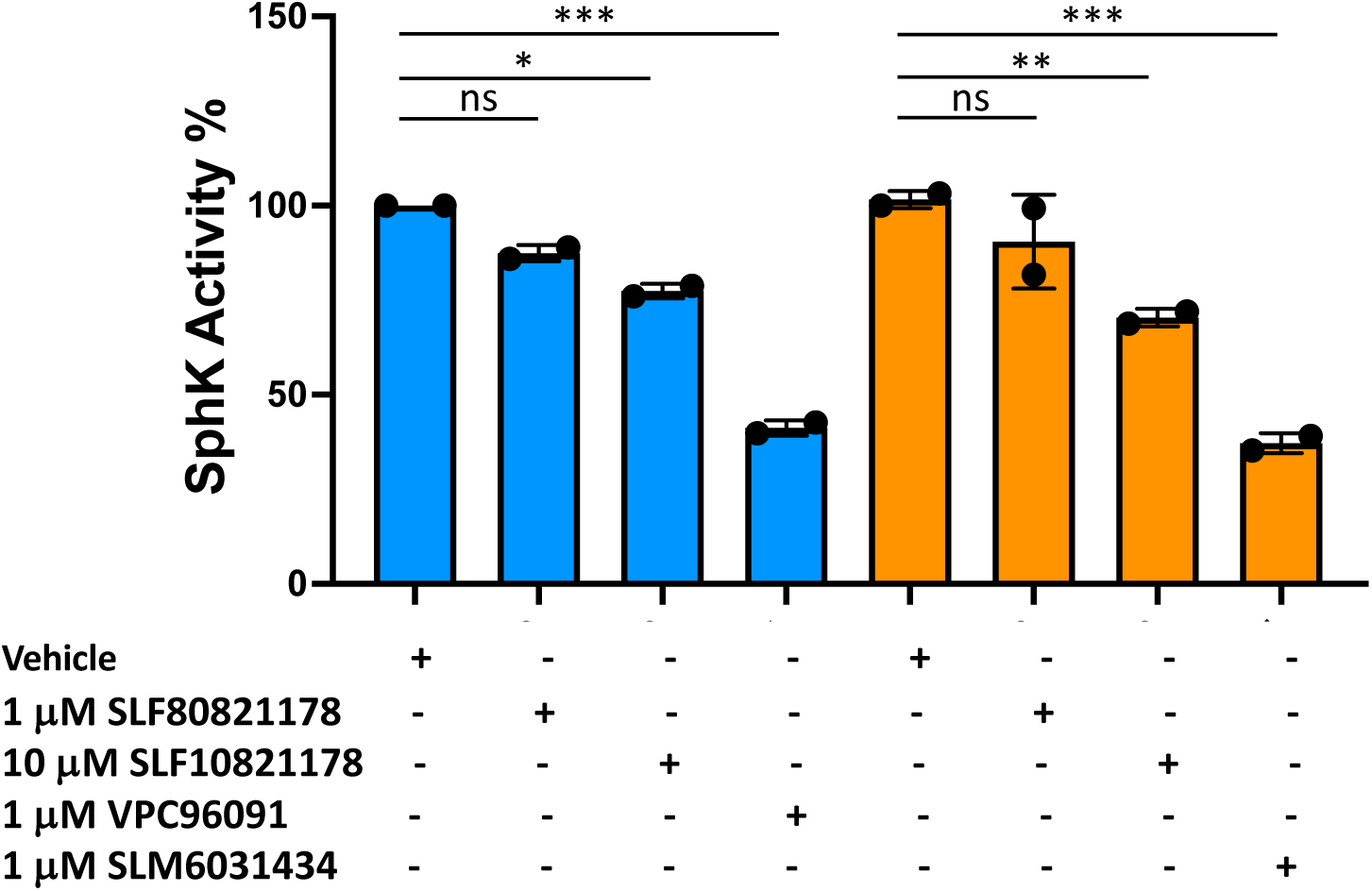
SLF80821178 counter-screens. **A**) Effect of SLF80821178 on release of S1P from C57BL/6 strain mouse red blood cells *ex vivo*. Control compound VPC96091 is a mouse sphingosine kinase type 1 (SphK1) inhibitor^49^. **B**) Effect of SLF80821178 on migration of C57BL/6 strain mouse splenocyte T lymphocytes. **C**) Inhibition of recombinant SphK1 (blue) and SphK2 (orange) by SLF80821178. Control compound SLM6031434 is an SphK2 inhibitor^50^.

### Efficacy in the MOG_35-55_ EAE model

Five SRMs (fingolimod, siponimod, ozanimod, ponesimod, etrasimod) are approved for treatment of various forms of multiple sclerosis (MS) and/or moderate-to-severely active ulcerative colitis. To learn whether STBs recapitulate the efficacy of SRMs in the standard preclinical model of MS (EAE – Experimental Autoimmune Encephalomyelitis) ^22–26^, we tested SLF80821178 in the C57BL/6 active EAE mouse model. As depicted in Fig. 3, administration of SLF80821178 before onset of symptoms (day 10) resulted in a significant decrease in peak clinical score (days 15-19) as well as a decrease in the chronic clinical scores (days 25-28) when compared to mice receiving only vehicle treatment. The efficacy of SLF80821178 is similar to the clinical course described in mice made deficient in Spns2 using inducible, whole body Cre recombinases on a *Spns2^fl/fl^* background^19,27^.

**Figure 3:**
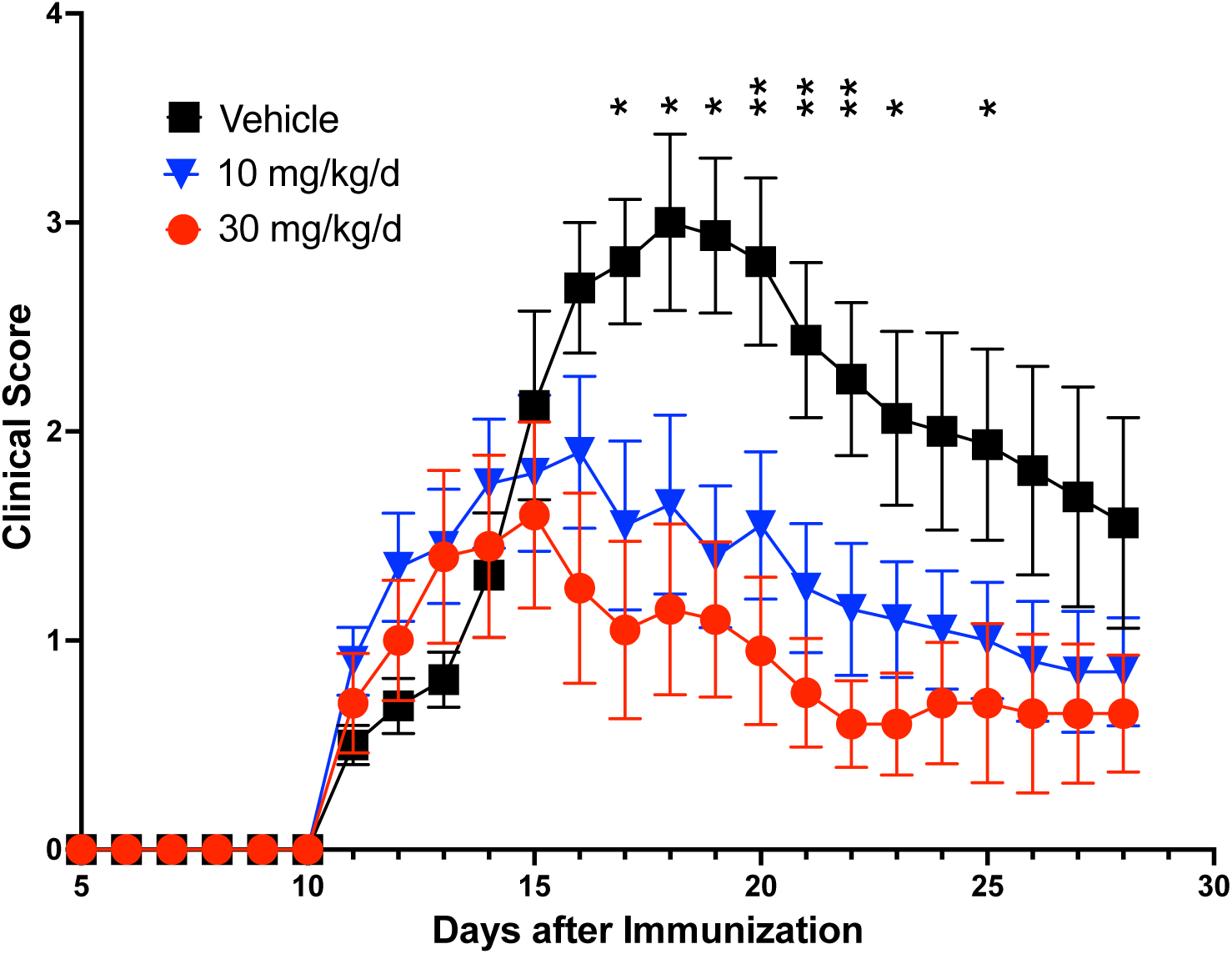
SLF80821178 efficacy in an EAE model. C57BL/6J strain female mice (n=10 /group) were vaccinated with MOG35-55 peptide and injected with pertussis toxin on day 0. On the 10^th^ day post-vaccination, the mice (all pre-symptomatic) were treated daily with vehicle or the indicated doses of vehicle or SLF80821178 (q.d., p.o.) for 14 days. Mice were scored by two individuals who were blinded as to the treatment regimen. Data are presented as mean ± SEM using the Kruskal-Wallis test. Stars indicate statistical significance when vehicle is compared to 30 mg/kg/d SLF80821178. Vehicle was 36.1 PEG400: 13 70% ethanol: 51.9 water (v/v) containing 4.6% (w/v) solutol.

### Cardiac toxicity

An on-target adverse effect of the SRM drug class is a transient bradycardia on starting therapy. This vagomimetic effect presumably results from agonist engagement of S1P1 receptors on the sinoatrial node cells in the heart^28^ (in rodents, this effect is mediated by the S1P3 receptor^29^). To learn whether an STB decreases heart rate on acute dosing, we administered SLF80821178 (100 mg/kg) or FTY720 (10 mg/kg) by oral gavage to freely behaving, instrumented rats and recorded heart rate (HR) and mean arterial pressure (MAP) by telemetry for 6 hours post dosing. SLF80821178 did not significantly perturb either parameter while fingolimod (FTY720) evoked the expected decrease in heart rate^28,29^ as well as an increase in MAP (Fig. 4).

**Figure 4:**
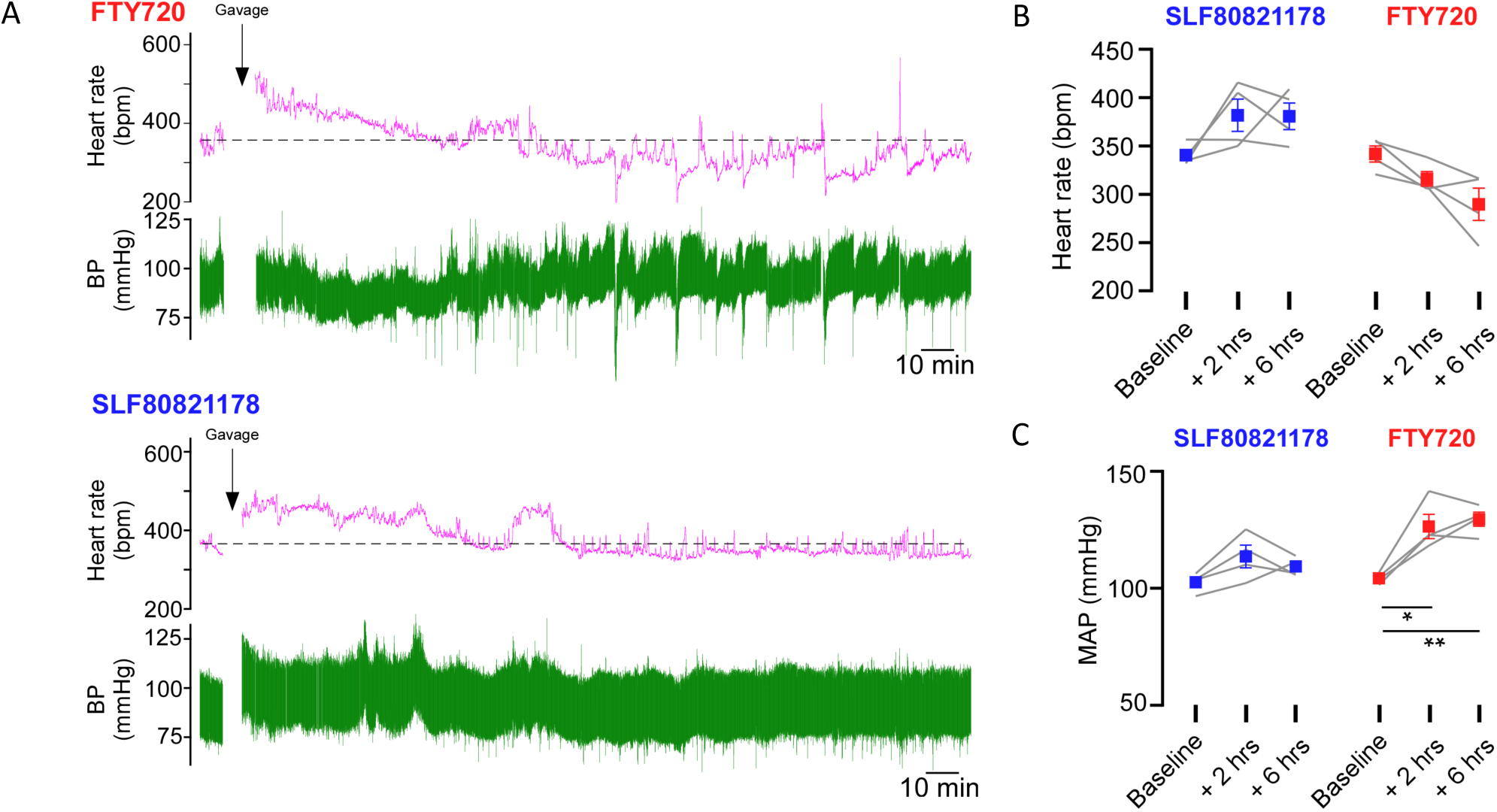
Cardiovascular effects of administration of SLF80821178 and fingolimod in rats. **A)** Experimental recordings of blood pressure and heart rate following administration (p.o.) of a single dose of SLF80821178 (100 mg/kg) or FTY720 (fingolimod) (10 mg/kg). **B)** Grouped data (mean ± SEM) for the effect of SLF80821178 and FTY720 on heart rate. SLF80821178 did not significantly affect mean heart rate (F_1.971, 5.913_ = 2.79, P=0.14), whereas FTY720 resulted in a significant reduction in heart rate (F_1.392,4.177_ = 8.91, P=0.035). **C)** Grouped data (mean ± SEM) for the effect of SLF80821178 and FTY720 on mean arterial blood pressure (MAP). SLF80821178 did not significantly affect MAP (mean arterial pressure) (F_1.390, 4.169_ = 4.528, P=0.095), whereas FTY720 resulted in a significant increase in MAP (F_1.308, 3.923_ = 27.23, P=0.0060). The effect of SLF80821178 and FTY720 on heart rate and MAP over time were assessed separately using repeated measures one-way ANOVA with Geisser-Greenhouse correction. * p<0.05 and ** p<0.01 based on Dunnett’s multiple comparisons test.

### Pulmonary toxicity

Another liability of the SRM drug class is the propensity of S1P1 receptor agonists to perturb endothelial barrier function, particularly at high drug concentrations^30^. For example, fingolimod administration is reported to significantly increase protein concentration in bronchoalveolar lavage (BAL) fluid in rats^31^. To discern whether this phenomenon occurred in response to an STB, we measured protein concentration in BAL fluid recovered from mice following SLF80821178, fingolimod, etrasimod or vehicle administration. We found that while there was no significant increase in BAL fluid protein in response to SLF80821178 compared to vehicle treated mice, both SRMs increased BAL fluid protein compared to vehicle treated animals (Fig. 5).

**Figure 5:**
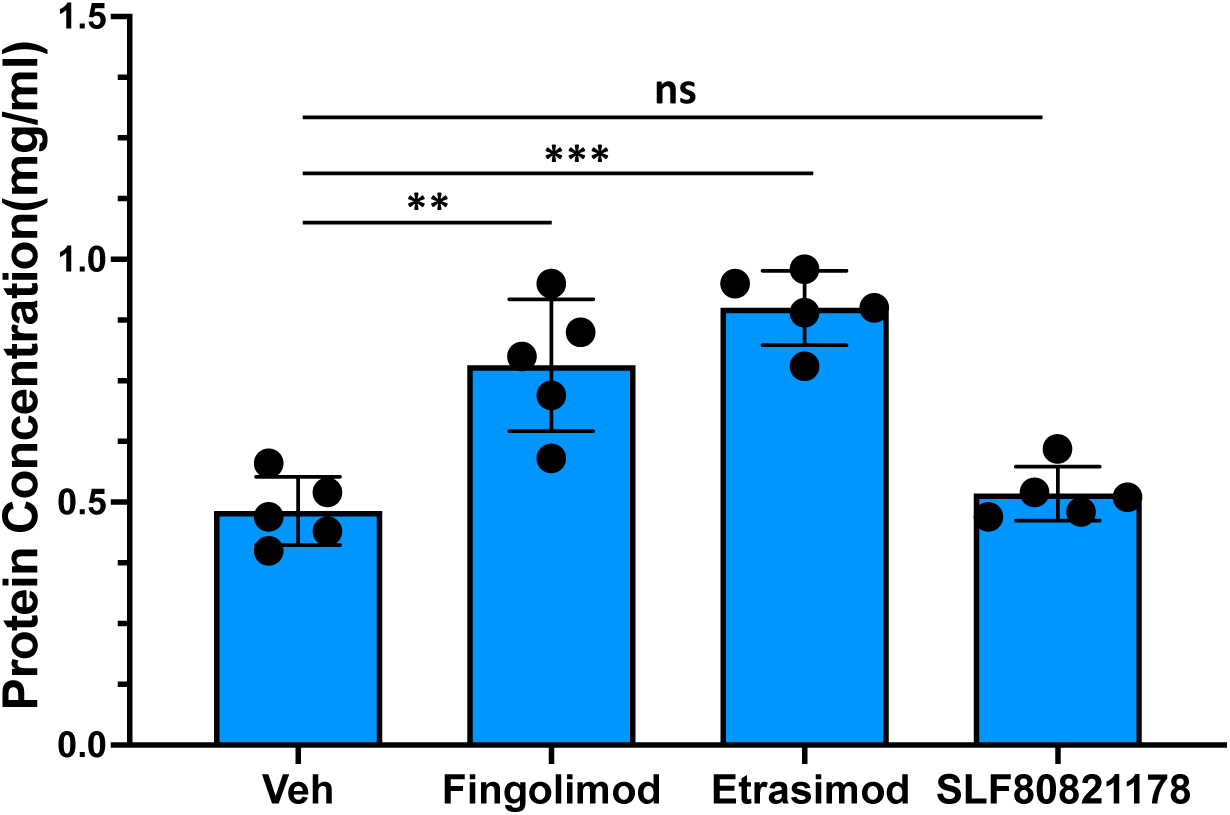
Effect of STB and SRM administration on protein content of BAL fluid. Cohorts of C57BL/6 strain mice (n = 5) were dosed (i.p.) with vehicle (10% hydroxypropyl-beta-cyclodextrin), fingolimod (3 mg/kg), etrasimod (5 mg/kg) or SLF80821178 (10 mg/kg). After approximately 4 hours, mice were euthanized, the trachea exposed and lungs were lavaged with 1 mL saline. Protein content in recovered BAL fluid (approximately 0.7 mL) was determined.

### Ototoxicity

Germ line disruption of either of two mouse genes encoding S1P pathway proteins – the S1P2 receptor and Spns2 – is associated with sensorineural hearing loss^32,33^. Spns2 deficiency has also been reported to be associated with human deafness^34^. To ascertain whether administration of an STB affects hearing acuity in adult mice, we administered SLF80821178 (20 mg/kg/d, i.p.) or vehicle chronically to cohorts of mice and tested the animal’s hearing on day 14 using ABR (Auditory Brainstem Response). As documented in Fig. 6, there is no discernable difference in hearing threshold between vehicle and SLF80821178-treated mice of either sex across a range of frequencies.

**Figure 6:**
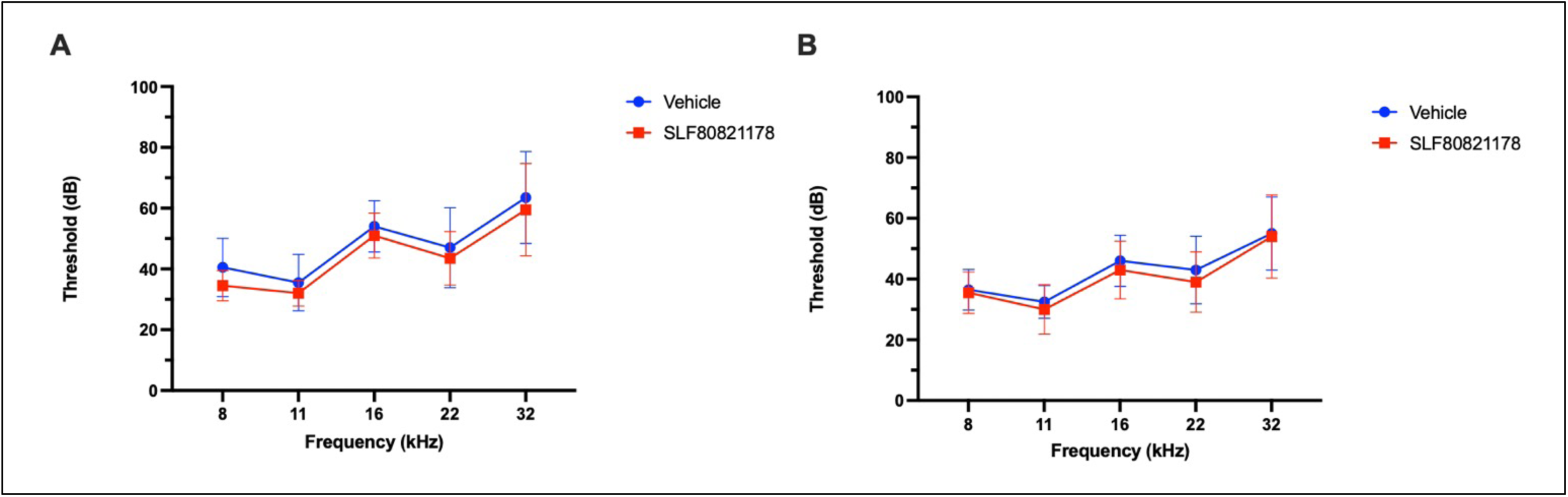
Effect of SLF80821178 on ABR. Auditory brainstem response (ABR) thresholds for cohorts of male (**A**) and female (**B**) mice treated for 14 days with SLF80821178 (20 mg/kg/d, i.p. (red) and vehicle control (blue). Graphs depict mean and error bars show standard deviation, N=10 mice per group. For both sexes there are no significant differences in ABR threshold at any frequency tested between SLF80821178 treated and vehicle treated mice (MWU tests of means at each frequency, p>0.05).

### Pharmacodynamics of STB administration

Ablation of *Spns2* in mice, either in the germ line^15,16^ or in the endothelial cell lineage *in utero*^17,18^, results in mouse strains with ∼50% decrease in blood lymphocyte counts compared to wild type littermates. Further, lymph S1P concentration, which was measured in some mouse strains, is nearly undetectable in the Spns2 deficient mice^17–19^. While the lymphopenia is consistently observed across multiple genetically manipulated mouse strains, the effect of *Spns2* deficiency on plasma S1P concentrations is variously reported to be either not changed significantly^15,19,34^ or decreased by up to 45%^16,17,35^. To this compendium, we add data obtained with our independently derived *Spns2*^-/-^ mouse strain, which was generated on a mixed B6-SJL strain background using Crispr/Cas9 technology. Our Spns2 null mice have ∼50% decrease in peripheral blood lymphocyte counts compared to *Spns2*^+/-^ or *Spns2*^+/+^ littermates (Fig. 7A), which is consistent with published studies. In agreement with some, but not other, previous reports our mouse strain had no significant difference in plasma S1P concentration among mice of different Spns2 genotypes (Fig. 7B). Finally, our *Spns2*^-/-^ mice had negligible thoracic duct lymph S1P concentrations compared to wild type or heterozygous mice (Fig. 7C), which is consistent with the existing literature^17–19^.

**Figure 7:**
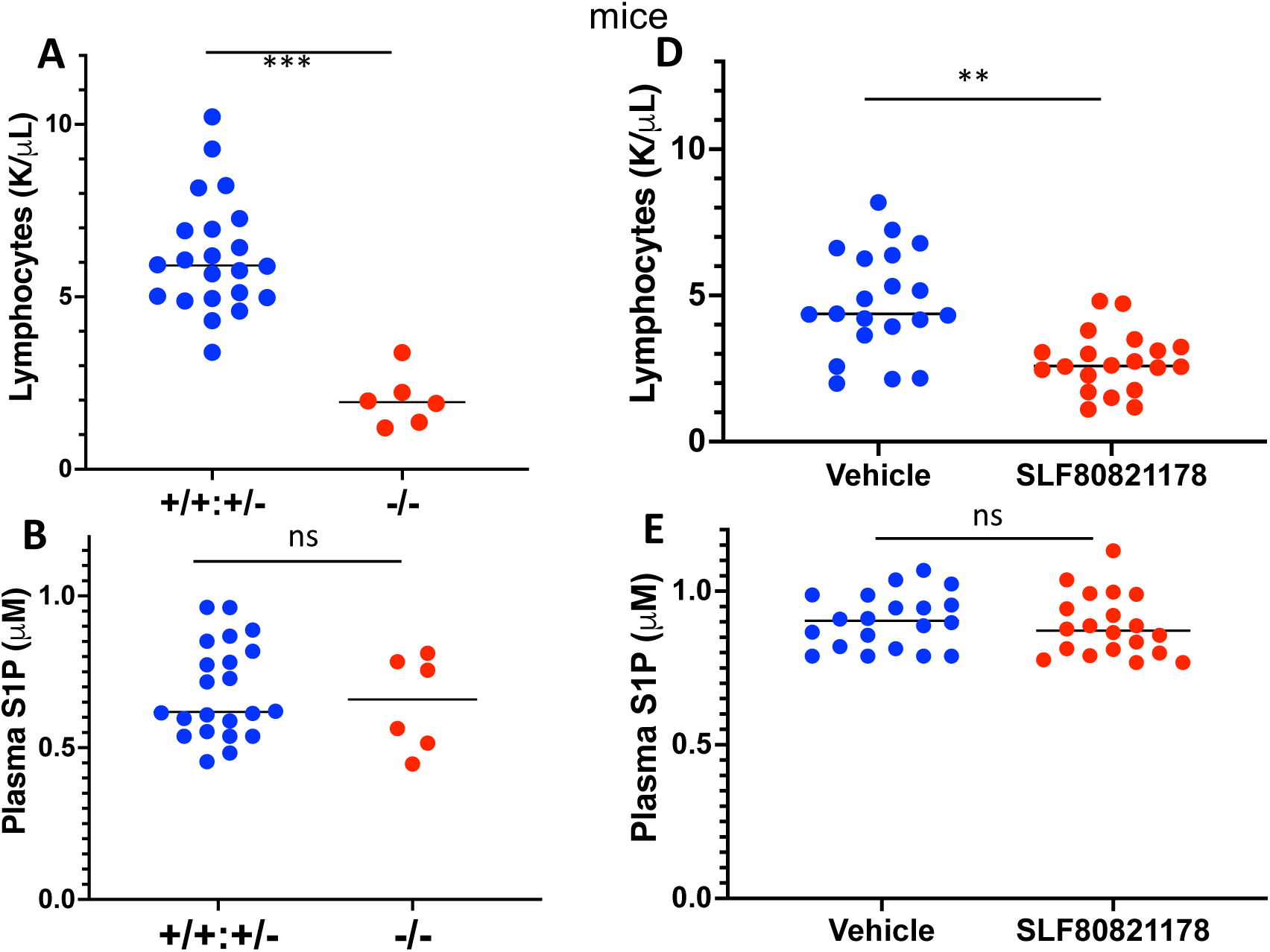
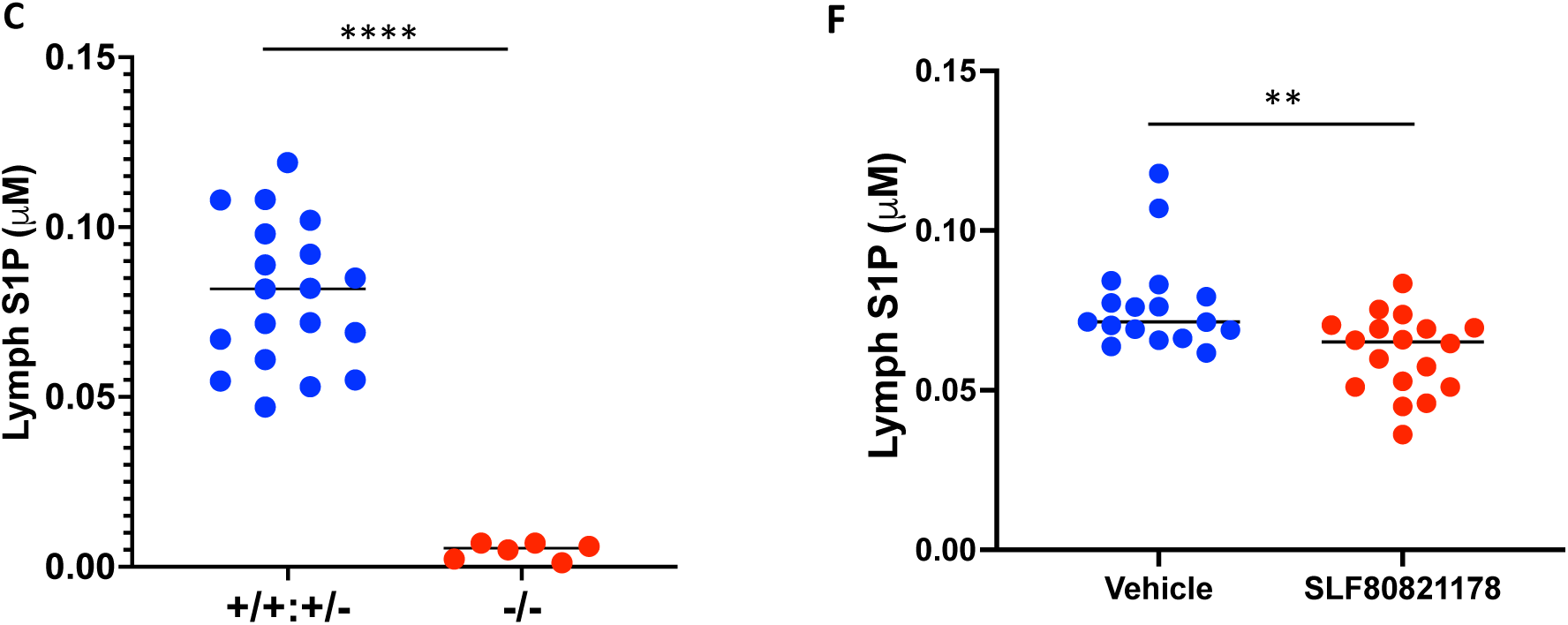
Pharmacodynamic markers in Spns2-deficient and SLF80821178 treated mice. **A - C**) *Spns2^-/-^* mice on a mixed SJL-C57BL/6 strain background were generated with Crispr/Cas9 technology (see Methods for details) and backcrossed for one generation with C57BL/6J mice. The resulting heterozygous mice were intercrossed and at 6-8 weeks of age were bled, dosed with mineral oil (o.1 mL, p.o.) and after one hr euthanized and lymph collected from the cisterna chyli. +/+ (wild type *Spns2*^+/+^) +/- (heterozygous *Spns2*^+/-^), -/- (homozygous (null) *Spns2*^-/-^). **D-F**) C57BL/6J mice of both sexes (6-8 weeks of age) were injected with vehicle or SLF80821178 (20 mg/kg/d, 15 days, i.p.). Two hrs after the final dose, mice were bled, dosed with mineral oil and after an additional 1 hr euthanized for lymph collection. **A,D**) peripheral blood lymphocyte counts, **B,E**) plasma S1P concentration, **C,F**) lymph S1P concentration. ** p < 0.01, *** p < 0.005, **** p < 0.001

The lymphopenia phenotype is recapitulated by STB administration. For example, single doses of SLF1081851^20^, SLB1122168^21^ and SLF80821178 (Foster *et al*., submitted) evoke dose-dependent decreases in total blood lymphocytes in C57BL/6 mice and Sprague-Dawley rats. Further, while plasma S1P concentrations were reduced ∼15% in response to our prototype STB, SLF1081851^20^, plasma S1P concentrations were unchanged in response to SLB1122168^21^ and SLF80821178 (Foster *et al*., submitted) after administration of a single dose of these inhibitors. Chronic dosing of SLF80821178 in C57BL/6J mice likewise evoked lymphopenia as measured on day 15 (Fig. 7D) without affecting plasma S1P concentrations (Fig. 7E).

In contrast to mouse strains wherein *Spns2* was ablated *in utero*, chronic SLF80821178 treatment of mice had minimal effect on lymph S1P concentrations (Fig. 7F). Likewise, acute dosing of SLF1081851, SLF1122168 or SLF80821178, while decreasing circulating lymphocyte counts substantially (Fig 8A), had no significant effect on lymph S1P concentration (Fig. 8B). Two explanations for this anomaly are either lymph endothelial cells are insensitive to SLF80821178 or the compounds do not accumulate in lymph. To investigate these possibilities, we determined the potency of SLF80821178 in blocking S1P release from cultured lymph endothelial cells (LECs) and measured S1P concentrations in lymph from mice dosed with SLF80821178. The potency of SLF80821178 in blocking S1P release from LECs (IC_50_ 23nM (Fig. 9A)) was similar to that measured with human monocytic leukemia U937 cells (IC_50_ 15nM (Fig. 9B)) and HeLa cells expressing mouse Spns2 (IC_50_ 53nM (Foster *et al*., submitted)). Further, S1P concentrations measured in lymph drawn 6 hours after administration of SLF80821178 were 4-5 times higher than in plasma from these animals (Fig. 9C, 9D).

**Figure 8:**
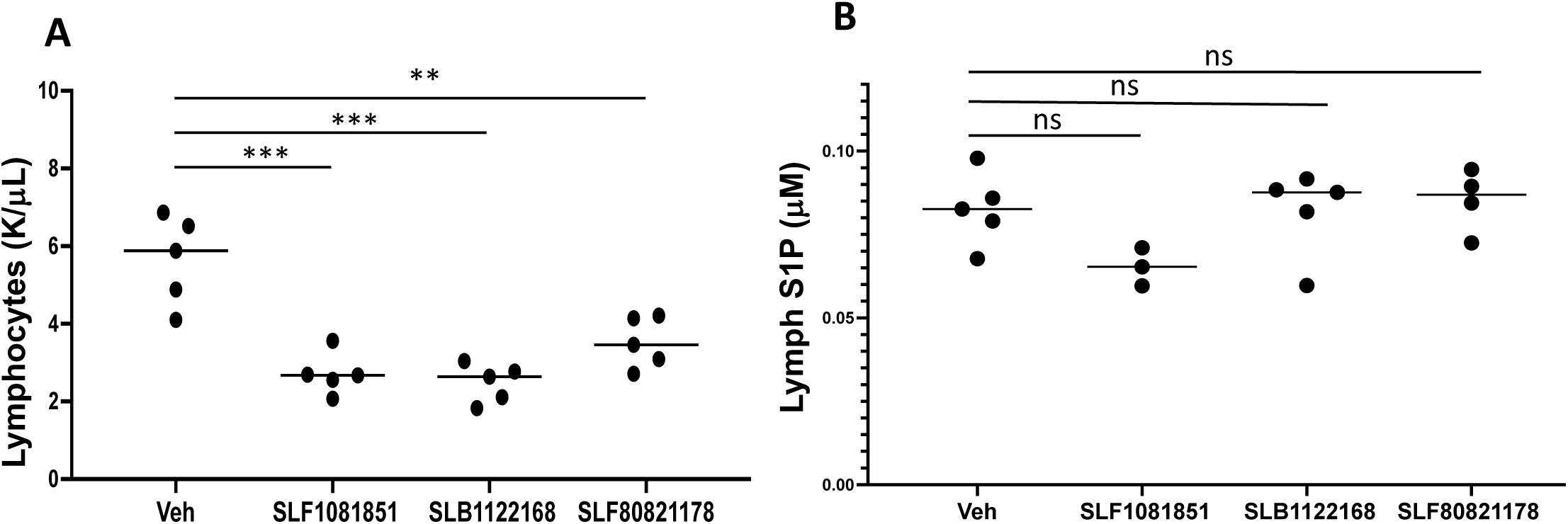
Effect of acute dosing of STBs on blood lymphocyte counts and lymph S1P concentrations. Cohorts (n=5 mice/group) of C57BL/6 mice were dosed with SLF1081851, SLB1122168 or SLF80821178 (20 mg/kg, i.p.). Two hrs after compound administration, mice were bled, dosed with mineral oil (0.1 mL, p.o.) and after an additional 1 hr euthanized for lymph collection (lymph collection failed in 3 mice). **A**) peripheral blood lymphocyte counts, **B**) lymph S1P concentrations.

**Figure 9:**
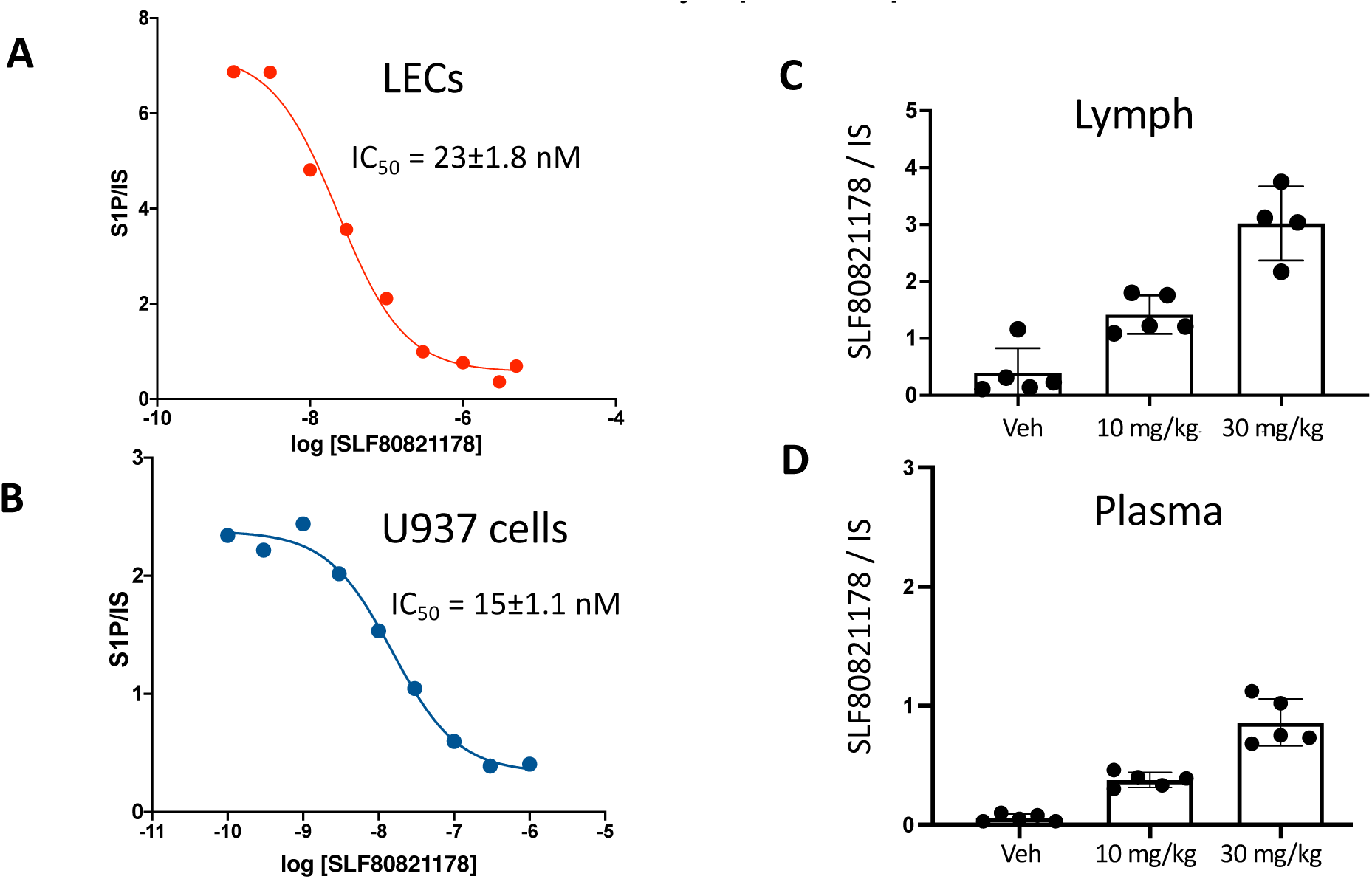
SLF80821178 inhibition of S1P release from cultured lymph endothelial cells (LECs) and presence in lymph. Cultured (passage 5-6) LECs obtained from adult human dermal tissue were plated onto a single 12-well plate in tested with the indicated concentrations of SLF80821178 (**A**). Cultured human monocytic U937 cells were placed in T-25 flask with indicated concentrations of SLF80821178 (**B**). See Methods for details of S1P release assay. Relative S1P levels in lymph (**C**) and plasma (**D**) drawn from female C57BL/6 mice 4 hrs (plasma) and 6 hrs (lymph) after administration of indicated doses of SLF80821178.

An additional point of differentiation between STBs and SRMs is the degree to which peripheral blood lymphocyte counts are suppressed in response to these agents, with the SRMs exerting an approximately 2-fold greater decrease than STBs. To investigate this phenomenon further, we used flow cytometry to discern the effect of a single dose of SLF80821178 or FTY720 (fingolimod) on subpopulations of circulating lymphocytes in mice. As documented in Fig. 10, both compounds reduce populations of B cells, NK cells and T cells but the SRM fingolimod has a proportionally greater effect of circulating T lymphocytes, particularly CD4+ positive cells, than does the STB SLF80821178. We note also that FTY720 has a disproportionally larger effect on blood double negative (CD4^-^, CD8^-^) T cells than SLF80821178. This population could warrant further characterization to better understand its physiological effect and genetic expression profile in relation to S1P biology.

**Figure 10:**
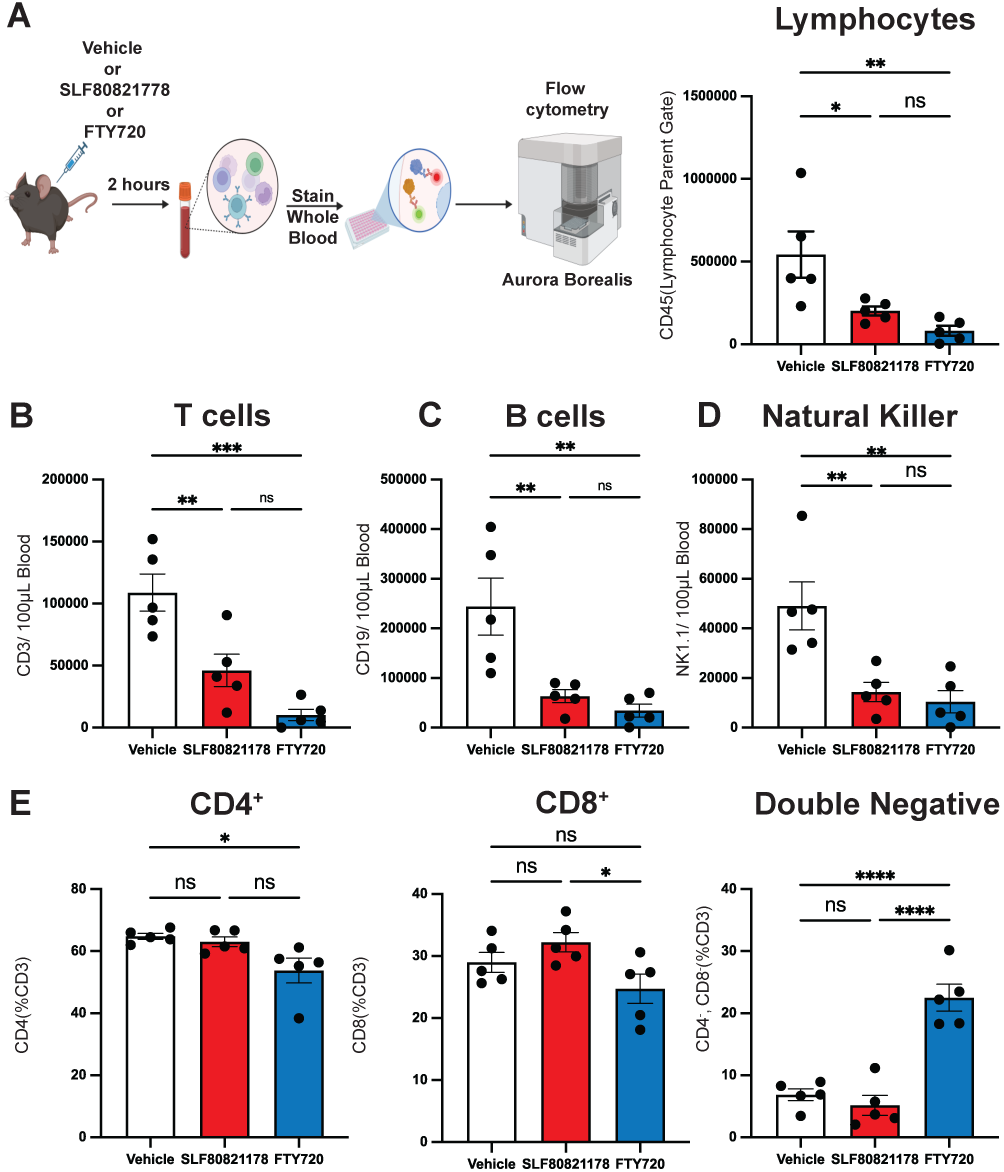
Impact of an acute dose of SLF80821178 or FTY720 on circulating lymphocyte populations. **A)** Schematic of experimental design. Flow cytometric analysis CD45^+^ cells from gated live lymphocytes isolated from whole blood of C57BL/6J mice 2-3 hours after administration of SLF80821178 (20 mg/kg, i.p.), FTY720 (3 mg/kg, i.p.) or vehicle control as identified by flow cytometric analysis (all groups N=5). Flow cytometric analysis of **B)** T-cells (CD3^+^), **C)** B-cells (CD19^+^), **D)** natural killer cells (NK1.1^+^). **E)** Flow cytometric analysis of CD4^+^ or CD8^+^ or double negative (CD4^-^CD8^-^) T-cells as a percentage of CD3^+^ cells. Data presented as mean ± SEM. Significance determined using One-way ANOVA with Sidak’s multiple comparisons test (A-E). * p < 0.05, ** p < 0.01, *** p < 0.001, **** p < 0.0001, ns = not significant.

## Discussion

We have found that the STB SLF80821178 is efficacious in a standard EAE model and that administration of SLF80821178 to mice evokes a significant decrease in total blood lymphocyte counts but had a minimal effect on the S1P concentrations of plasma and lymph. We found further that unlike SRMs, SLF80821178 neither decreased heart rate nor compromised the lung vascular barrier. STBs and SRMs differ also in that the former suppresses blood lymphocyte counts to a lesser extent than the latter (ca. 45% vs. 90%), which is reflected by the relatively lesser effect of an STB on peripheral T lymphocyte populations. While it is logical that less lymphopenia equates to less immunosuppression, we do not know whether the extent of lymphopenia in response to these drugs is proportional to the extent to which they immunocompromise an animal. Finally, chronic dosing of SLF80821178 did not decrease hearing acuity in adult mice. Because a neurosensorial hearing defect is a phenotype associated with *Spns2*^-/-^ mice^33^, it is particularly important to test and STB for ototoxicity as such a signal in a preclinical animal model would substantially decrease enthusiasm for Spns2 as a drug target. A comparison of the effects of STBs, SRMs and disruption of *Spns2* in the mouse embryo is summarized in Table 1.

**Table 1:**
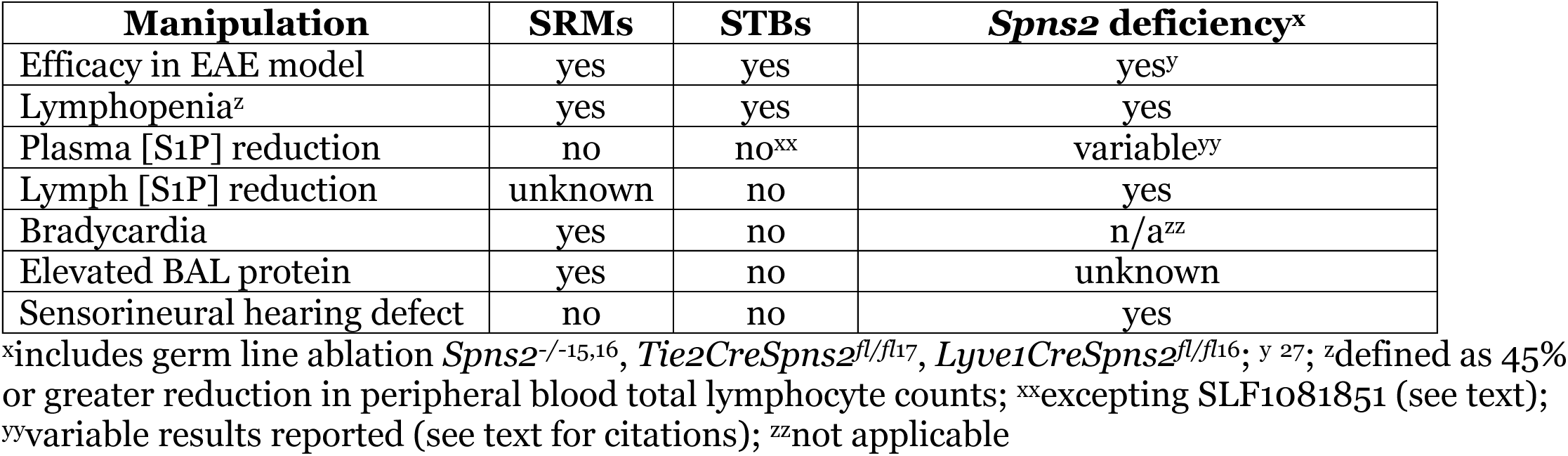
Comparison of *in vivo* effects of acute S1P transport (Spns2) blockade (STB), S1P receptor modulation (SRM) and Spns2 deficiency achieved via genetic manipulations of mice.

| Manipulation | SRMs | STBs | *Spns2* deficiency<sup>x</sup> |
| --- | --- | --- | --- |
| Efficacy in EAE model | yes | yes | yes<sup>y</sup> |
| Lymphopenia<sup>z</sup> | yes | yes | yes |
| Plasma [S1P] reduction | no | no<sup>xx</sup> | variable<sup>yy</sup> |
| Lymph [S1P] reduction | unknown | no | yes |
| Bradycardia | yes | no | n/a<sup>zz</sup> |
| Elevated BAL protein | yes | no | unknown |
| Sensorineural hearing defect | no | no | yes |

<sup>x</sup>includes germ line ablation *Spns2*<sup>-/-15,16</sup>, *Tie2CreSpns2*<sup>fl/fl17</sup>, *Lyve1CreSpns2*<sup>fl/fl16</sup>; <sup>y</sup> 27; <sup>z</sup>defined as 45% or greater reduction in peripheral blood total lymphocyte counts; <sup>xx</sup>excepting SLF1081851 (see text); <sup>yy</sup>variable results reported (see text for citations); <sup>zz</sup>not applicable

Our data, albeit limited in scope, support the proposition that STBs recapitulate the efficacy of SRMs with less toxicity. The shared efficacy in the EAE model is not surprising in that both STBs and SRMs depress circulating lymphocyte numbers and mice rendered deficient in Spns2 by inducible Cre recombinase (*Mx1Cre*;*UbcCre^ERT^;Spns2^fl/fl^*) are protected in the EAE model^19,26^. Further, because STBs neither affect circulating S1P tone nor engage S1P1 receptors, it is not surprising that we observed no perturbation of either heart rate or the pulmonary vascular barrier in response to SLF80821178. The observed hearing deficiency in Spns2 null mice indicates a requirement for Spns2 for proper development of the inner ear, but the hearing phenotype generated by genetic manipulation of the germ line is apparently not predictive of the drug’s action in adult animals. We do not understand the discrepancy among literature reports regarding plasma S1P concentrations in response to Spns2 deficiency or why our prototype STB, but not later generations STBs, reduces plasma S1P concentrations slightly. However, the practical value of our observation is that plasma S1P concentration is not a useful biomarker of STB target engagement. In contrast, the sum of our experience with STBs leads us to suggest that, like SRMs, peripheral blood lymphocyte counts are a reliable pharmacodynamic index.

Spns2 expression during mouse embryogenesis is necessary for normal thoracic duct lymph S1P concentrations (100-200 nM) as judged by examination of lymph drawn from the cisterna chyli of adult mice^16–19^ (and this report). However, we observed only a minimal effect of STB administration on this parameter despite the high potency of SLF80821178 in blocking the release of S1P from cultured lymph endothelial cells. Further, STBs such as SLF80821178 accumulate in lymph. These observations discount the explanation that lack of drug contact or response by lymph endothelium is responsible for our observations. Unfortunately, we are unable to proffer an explanation for this anomaly, perhaps due to our limited understanding of either STB action or S1P transport biochemistry. Existing knowledge of Spns2 (or Mfsd2b) function is largely informed by phenotypes observed when the underlying genes are inactivated in embryonic animals, *i.e*. zebrafish^36,37^ and mice (*vide infra*). As evidenced in this and numerous other reports, the predictive value of results regarding drug action from genetic manipulation is limited. Further, our reliance on phenotypic assays for STB characterization precludes certainty regarding the target contacted by the inhibitors although recently reported cryoEM structures of human Spns2^38–41^ with SLF1081851 (16d) or SLB1122168 (33p) bound indicates that the proximal STB target is Spns2.

A convenient explanation of Spns2 function has this exporter releasing S1P from lymph endothelial cells to lymph and therefore predicts that STB administration would suppress lymph S1P concentrations, flatten the lymph: lymph node S1P concentration gradients and thereby inhibit lymphocyte egress to efferent lymph. The negligible effect of the three STBs tested on bulk lymph S1P concentration when administered at doses that drive lymphopenia indicates that this narrative is incorrect. Thus, the question ‘How does interfering with Spns2-dependent S1P transport disrupt lymphocyte trafficking?’ remains unanswered. Garcia-Seyda *et al.* reported^42^ that S1P is not a long range chemoattractant of naïve human T lymphocytes and they proposed rather that it is either a narrow S1P gradient or a stromal gating effect at the lymph node paracortical sinus endothelium that underlies the dependence of T cells on S1P for egress to medullary sinuses and thereby on to efferent lymph. The roles of the S1P phosphatase, Plpp3 (LPP3), and Spns2 in the S1P-dependent eggress of mature thymocytes from thymus to blood reported by Bréart *et al*.^3^ likewise suggests a narrow gradient / stromal gating effect. Indeed, Wei *et al*.^43^, on the basis of data obtained by intravital microscopy of lymph nodes, suggeted that a similar stromal gating effect might also explain the interdiction of lymphocyte eggress by the SRM fingolimod. The stromal gating and narrow S1P concentration gradients models are both consistent with our observation that acute dosing of STBs drives lymphopenia without changing lymph S1P concentrations significantly.

### Limitations of this study

The salient limitations of this study are that we report STB efficacy in only a single disease model (EAE) and we relied on a single compound, SLF80821178, for most of the results reported. Additional testing of more STBs in other autoimmune disease models as well as in allograft models (wherein SRMs are notably effective) is necessary to test fully our hypothesis that SRMs and STBs have comparable efficacies. Regarding our reliance on SLF80821178 for these studies, our results with this compound are entirely consistent with those we have obtained with STBs on other chemical scaffolds excepting a modest effect on plasma S1P concentrations (15% decrease), where our prototype STB, SLF1081851, is an outlier. Additionally, our measurements of lymph S1P concentrations are restricted for technical reasons to lymph drawn from the caudal aspect of the thoracic lymph duct (cisterna chyli), where many lymphatic vessels (including lacteals) terminate. Lymph S1P concentrations in the efferent lymph ducts of, for example, individual lymph nodes might be different from those in bulk lymph. Likewise, the sensitivity of S1P release to SLF80821178 exhibit by cultured human LECs might not fully recapitulate that of mouse endothelium.

## Materials & Methods

### RBC S1P release assay

A mouse was exsanguinated under urethane anesthesia using EDTA as an anti-coagulant. The blood was centrifuged at 1,600 x g for 10 min at 4°C. After discarding plasma and removing the buffy coat (leukocytes), the RBCs were washed twice with 1 mL cold PBS and centrifuged at 300 x g for 10 min at 4°C. The washed RBCs were resuspended in 1 mL of cold PBS and counted with a blood analyzer (Element HT-5, Heska, Loveland, CO). RBCs (30 µL, 100-120 x 10^6^) were added to a 2 mL microcentrifuge tube containing 1 mL Tyrodes buffer supplemented with 1 mM glucose, 0.2% fatty acid-free BSA and 1 µM C17-sphingosine. After incubation in a Thermoshaker (30 min, 37°C, 400 rpm), RBCs were pelleted by centrifugation (1,600 x g for 10 min) and an aliquot of the supernatant fluid prepared for quantifying C17-S1P by LCMS as described for plasma extraction (see below).

### Splenocyte isolation

A fresh whole mouse (C57BL/6v strain) spleen was excised following euthanasia under urethane anesthesia and minced into small pieces in Hank’s balanced salt solution (HBBS). The pieces were pressed through a cell strainer (70 µm nylon mesh, Fisher Scientific) using a syringe plunger and collected into a 50 mL conical tube, washing through the strainer with HBBS. The cell suspension was centrifuged at 300 x g for 5 minutes at room temperature and the supernatant fluid was discarded. The cell pellet was resuspended in 500 µL HBBS with 0.5% fatty acid-free BSA and lymphocytes were isolated by using the EasySep Mouse Streptavidin RapidSpheres isolation kit (STEMCELL Technologies, Vancouver, British Columbia, Canada) with the protocol provided. Briefly, 10 µL FcR blocker solution, 5 µL Biotin Rat Anti-Mouse TER119 and 5 µL Biotin Rat Anti-Mouse CD45R (BD Biosciences, Franklin Lakes, NJ) were added into the sample. The mixture was incubated at room temperature for 10 minutes, followed by adding 50 µL magnetic streptavidin beads and incubating for 2.5 minutes. The mixture was diluted to 2 mL with PBS containing 0.5% fatty acid-free BSA and over 2.5 minutes bound cells were immobilized with a magnet. The cell suspension was transferred to a new tube and centrifuged at 300 x g for 5 minutes to pellet the cells, which were resuspended in the RPMI 1640 media with 0.5% BSA (fatty acid free) at 2.5 x 10^6^ cells / mL for the migration assay.

### Splenocyte Migration Assay

After serum starvation in RPMI 1640 medium supplemented with 0.5% fatty acid-free BSA for 3 h at 37°C, lymphocytes (2.5 × 10^5^ cells) in 100 µL of media were added to the upper wells of 24-well tissue culture inserts with a 3 micron polycarbonate membrane (Costar) in, with 600 µL or media supplemented with S1P (100 nM), SLF80821178 (0.1 or 1 µM) dilution or medium in the bottom wells. The assay was conducted at 37°C in 5% CO2 for overnight. Three wells were set up for each experimental conditions and combined for centrifugation to concentrate the migrated cells into 60 µL. The migrated cells were counted with a TC20 automated cell counter (Bio-Rad, Hercules, CA).

### Sphingosine kinase (SphK) assay

Mouse SphK1 and SphK2 inhibitory activity of SLF80821178 was assessed using a previously published method^44^. Briefly, cell lysates were prepared from recombinant baculovirus infected Sf9 cells in 20 mM Tris-Cl (pH 7.4), 1 mM 2-mercaptoethanol, 1 mM EDTA, 5 mM sodium orthovanadate, 40 mM beta-glycerophosphate, 15 mM NaF, 1 mM phenylmethylsulfonyl fluoride, 10 mM MgCl2, 0.5 mM 4-deoxypyridoxine, 10% glycerol, and 0.1 mg/ml each leupeptin, aprotinin, and soybean trypsin inhibitor. SphKs assays were performed in presence of D-*erythro*-sphingosine, [γ-^33^P]ATP and SLFF80821178 at the concentrations indicated. Radiolabeled S1P was recovered by organic solvent extraction followed by thin-layer chromatography. Signals were localized by autoradiography and the analyte quantified by scintillation counting after scraping the tlc plate. All measurements were made in duplicate.

### EAE

Experimental autoimmune encephalomyelitis (EAE) was induced in female mice (10 weeks old) as described.^45^ The MOG35-55 peptide (100 µg, CSBio CS0681) was emulsified in complete Freund’s adjuvant containing *Mycobacterium tuberculosis* (1 mg/mL, Sigma F5881). *M. tuberculosis* (BD231141) was added to achieve a final concentration of 4 mg/mL. The peptide mixture was then injected subcutaneously (100 µL) at the base of the tail. Pertussis toxin (200 ng, List Biologics, Campbell, CA) was administered intraperitoneally on the day of MOG immunization and repeated 1 day after. The mice were assessed daily by two blinded evaluators using the following scoring system: 0 – no clinical disease, 1 – limp tail, 2 – hindlimb incoordination, 3 – hindlimb weakness, 4 –hindlimb paralysis, 5 – moribund. Compounds were administrated once daily by oral gavage starting at Day 10 and continuing for 14 days.

### BP recordings and analysis

Adult male Sprague-Dawley rats (N=4) were singly housed during experiments in standard rat cages (45cm x 30cm x 20 cm) under 12:12 light: dark cycle (lights on 7am) at 23°C-24°C with water and food provided *ad libitum*. Rats were anesthetized with 2.5% isoflurane in pure oxygen for implantation of a radiotelemetry probe (PA-C10; Data Sciences International) in the right femoral artery. Following 7 days’ recovery, pulsatile arterial pressure was measured in unanesthetized freely-behaving conditions. BP data was acquired at 500 Hz with a CED 1401 A/D acquisition system using Spike2 (Version 8 & 9, Cambridge Electronic Design, UK). SLF80821178 (100 mg/kg), fingolimod ((FTY720) catalog #11975, Cayman Chemical Co., Ann Arbor, MI) or vehicle was administered by oral gavage in randomized order during the light phase (0900-1200 hours) with at least 48 hrs. between drug administration. Mean arterial pressure (MAP) and heart rate (HR) were derived from pulsatile arterial pressure using a publicly available script (HRBP8 script from Cambridge Electronic Design, UK, www.ced.co.uk/downloads/scriptspkanal). For analysis, mean values of MAP and HR over a 30 s period were extracted in 30 minute intervals starting 2 hours before drug administration and continuing for 6 hours after drug administration. Rats were purchased from Inotiv (West Lafayette, IN).

### Bronchoalveolar fluid (BAL) collection

Four hours after administration of vehicle, fingolimod, etrasimod ((APD334) catalog #21661, Cayman Chemical Co., Ann Arbor, MI) or SLF80821178, mice were euthanized by isoflurane overexposure. An incision in the ventral neck skin was made to expose the trachea. A 26-gauge needle was used to puncture the trachea and a catheter was inserted about 0.5 cm. A sterile balanced salt solution (1 mL) was introduced into the lungs via the catheter; 0.6-0.7 mL of the solution was recovered from the lungs. Protein concentration in the BAL fluid was measured with the standard BCA method.

### Auditory brainstem response (ABR) measurement

ABRs of C57BL/6j mice were measured two weeks after the mice began receiving injections of either SLF80821178 (20 mg/kg/d, i.p., 14 days) or a vehicle control. Researchers were blinded to the treatment group of the mice during ABR acquisition. Mice were anesthetized with an intraperitoneal injection of 20 mg/kg ketamine xylazine hydrocholoride (Ketaset, Zoetis; AnaSed injection, Akorn Animal Health). ABRs were recorded as previously described^46^. Briefly, anesthetized mice were placed within a sound-attenuating chamber for ABR acquisition (Med-Associates, product number: ENV-022MD-WF). ABRs were obtained using TDT’s ABR acquisition system for small mammals (Tucker Davis Technologies, Alachua, FL). Recordings were conducted using subdermal needle electrodes (FE-7; Grass Technologies, West Warwick, RI). The inverting electrode was inserted over the mastoid of the right ear to detect signal from the auditory nerve. The noninverting electrode was placed along the nose of the mouse at the vertex of the midline, and the ground electrode was situated on the upper thigh. Pure tone stimuli were presented to mice at a rate of 21/sec via the RZ6 Multi I/O Processer (Tucker Davis Technologies, Alachua, FL). Responses were filtered from 300-3000 Hz and threshold levels were determined for stimuli presented at 8, 11, 16, 22, and 32 kHz. The intensity of the stimulus was decreased 5-10 dB at each frequency until a response waveform could not be identified, and each intensity was tested twice to confirm a lack of response. If a response waveform could not be identified at even the loudest intensity, the threshold was determined to be the maximum intensity plus 5 dB.

### Blood sampling

Mice were anesthetized to effect with isoflurane and 50-100 µL blood was drawn from the retro-orbital sinus using an EDTA coated micropipet and collected in a microcentrifuge tube containing 2 µL of 0.5 M EDTA. Lymphocytes were counted using a blood analyzer (Element HT5, Heska, Loveland, CO).

### Plasma preparation

Blood was centrifuged at 800 x g at room temperature for 10 min. The supernatant fluid was transferred to a fresh microcentrifuge tube and centrifuged at 10,000 x g for 10 min. The supernatant fluid (platelet-poor plasma) was prepared for LC-MS/MS analyses by adding 10 µL of plasma to 420 µL of a 0.2 % BSA (fatty acid free) solution containing internal standard (20 µL of 0.5 µM *d7*-S1P (2.5 pmoles)). Protein was precipitated by adding 50 µL of 100% trichloroacetic acid, mixing, incubating on ice for 30-45 min and collected by centrifugation (10,000 x g at 4°C). The pellet was washed once with 1 mL cold water, re-centrifuged (10,000 x g) at in room temperature. After discarding the supernatant fluid, the pellet was soaked in 0.3 mL methanol (Optima grade, Fisher Scientific) at room temperature for 15 minutes followed by centrifugation (12,500 x g, room temperature). An aliquot of the supernatant fluid (160 µL) was transferred to a vial insert for LCMS analysis.

### Lymph isolation

To recover lymph from mice, we used the protocol of protocol of Nagahashi *et al*.^47^ with minor modifications. Briefly, the animals were dosed by oral gavage with 0.1 mL of olive oil about one hour prior to euthanasia. At necropsy, the abdominal cavity was opened while avoiding bleeding and the intestinal trunk was gently moved aside to visualize the (white) cisterna chyli. The cisterna chyli was punctured with a 30 gauge beveled needle and 2-10 µL of lymph was collected into a micro pipet tip. Any samples with a pinkish tinge (indicating blood contamination) were discarded. Lymph was prepared for LCMS analysis as described for plasma.

### Sample preparation of biologic materials for LCMS

10 µL of blood or plasma or 2-10 µL of lymph were diluted into 420 µL of water containing 0.1% fatty acid-free BSA. Internal standard (*d7*-S1P, Avanti Polar Lipids, Alabaster, AL, 2.5 pmoles) was added and after vortex mixing, 50 µL of 100% trichloroacetic acid (w/v) in water was added followed by further mixing. Samples were held on ice for ∼45 minutes and centrifuged at 10,000 x g for 10 minutes in the cold room. The supernatant fluid was discarded and the pellets were washed once by vortex mixing with 1 mL of ice cold water. After further centrifugation, the supernatant fluid was removed by aspiration and discarded. The pellet was soaked in 300 µL of methanol at room temperature for 15 min. After a final centrifugation (11,000 x g, 5 min) at room temperature, about one-half of the supernatant fluid was transferred to an UPLC vial.

### LCMS Methodology

S1P, internal standard (*d7*-S1P), sphingosine and SLF80821178 quantification were performed using a tandem quadrupole mass spectrometer (Waters Xevo TQ-S micro) with a UPLC inlet (Waters Acquity h-class+) with a reverse phase C-18 column (Waters BEH C-18 1.7 µm bead size, 2.1mm x 50mm). Our chromatography protocol, which is a modification of that described by Frej *et al.*,^48^ was a binary solvent gradient with a constant flow rate of 0.4 mL/min with a column temperature of 60°C. Mobile phase A (MPA) consisted of water/methanol/formic acid (79:20:1) while Mobile phase B (MPB) was methanol/acetone/water/formic acid (68:29:2:1). Following injection (1-9 µL) on column, a run began with 50:50 MPA: MPB for 0.5 min followed by increasing MPB linearly to 100% to 3.5 min and holding at 100% MPB for another 3 min. The column is re-equilibrated to 50:50 MPA:MPB for 1.5 min. Analytes were detected using these MRM protocols: S1P (380.1>264.4, voltages: cone 18, collision 16), deuterated *d7*-S1P (387.2>271.4, voltages: cone 24, collision 16), sphingosine (300.3>252.3, voltages: cone 30, collision 18) and SLF80821178 (346.11>86.94 cone 2, collision 16), all in ESI+ mode. Peak analysis was accomplished using Waters TargetLynx software ver. 1.4. Solvents were LCMS Optima grade (Fisher Scientific, Waltham, MA). S1P concentrations were estimated from the ratio of S1P signal to internal standard (*d7*-S1P) signal.

### Mice

Generation of the *Spns2^-/-^* mouse strain: CRISPR gene editing technology was used to generate the *Spns2* mutant mouse strain. A gRNA targeting *Spns2* exon 3 was selected based on the search via the CRISPR guide design algorithm CRISPOR (http://crispor.tefor.net/). crRNA, tracrRNA and Cas9 were purchased from IDT (Coralville, Iowa). crRNA and tracrRNA were diluted to 100 µM in RNase-free microinjection buffer (10mM of Tris-HCl, pH 7.4, 0.25mM of EDTA). 5 µl crRNA and 5 µl tracrRNA were mixed and annealed in a thermocycler by heating the mixture to 95°C for 5 minutes and ramped down to 25°C at 5°C/min. Ribonucleic protein (RNP) complex was formed by mixing and incubating Cas9 at 0.2 µg/µl with crRNA/tracrRNA at 1 µM in RNase-free microinjection buffer at 37°C for 10 minutes. The fertilized eggs were collected from B6-SJL females mated with the males of the same strain. The RNP was delivered into the fertilized eggs by electroporation with a NEPA21 super electroporator (Nepa Gene Co., Ltd. Chiba, Japan) under the following conditions: 2 pulses at 40 V for 3 msec with 50 msec interval for poring phase; 3 pulses at 7 V for 50 msec with 50 msec interval for transferring phase. The treated zygotes were cultured overnight in KSOM medium (EMD Millipore, Billerica, MA) at 37°C in 5% CO2. The following morning, zygotes that had reached the two-cell stage were implanted into the oviducts of pseudopregnant foster mothers (ICR strain, Envigo, Indianapolis, IN). Pups born from the foster mothers were screened using tail snip DNA by PCR genotyping followed by Sanger’s sequencing. Germline transmission of the desired knockout alleles was confirmed by breeding the founders with wildtype C57BL/6j mice. Wildtype mice: Mice without genetic manipulation used in these studies were the C57BL/6j strain (purchased from Jackson Laboratories, Bar Harbor, ME) or the C57BL/6v strain (from our breeding colony that was started with C57BL/6j mice). All procedures with animals were approved by the University of Virginia Animal Care and Use Committee prior to experimentation.

### Culturing & Assaying LECs

Adult human dermal Lymphatic Endothelial Cells (LECs, PromoCell, catalog number C-12217) were cultured on fibronectin-coated plates in Endothelial Cell Growth Medium MV (PromoCell, C-22020). Nearly confluent LEC cultures at 5-6 passages were seeded onto 12-well cell culture plates (1 x 10^6^ cells/well) for the S1P release assay. S1P release from cultured LECs was measured as described for Spns2 plasmid DNA transfected HeLa cells and U-937 cells^8^. Briefly, LECs were cultured on 12 well plates until nearly confluent. To begin the assay, culture media was removed, the cell monolayers washed with PBS and the cells incubated in 2 mL of serum-free culture media supplemented with a cocktail of S1P degradation inhibitors (sodium fluoride, sodium orthovanadate, 4-deoxpyridoxine) and 0.2 % fatty acid free BSA. After 16-18 hours, 1.8 mL of media was transferred to a microfuge tube, 2.5 pmoles of internal standard (*d7*-S1P, Avanti Polar Lipids) was added and protein with bound S1P is concentrated by the addition of 0.2 mL 100% trichloroacetic acid. Following centrifugation of precipitated protein, pellets were washed with cold water, soaked in 0.3 mL LCMS grade methanol for 15 minutes and, after further centrifugation, an aliquot of the supernatant fluid was transferred to LC vials.

### Flow cytometry of whole blood leukocytes

Whole blood was collected by cardiac puncture from mice two hours after administration of SFL80821178 (20 mg/kg, i.p.) or FTY720 (2 mg/kg, i.p.) or vehicle control. Blood (100 µL) was treated with 10U of heparin in sterile saline. Splenocytes were isolated by grinding through a 100µm cell strainer and washing with PBS containing 2mM EDTA. Samples were then incubated with ACK lysis buffer for 2 minutes and centrifuged at 400 x g for 5 minutes. Fc-receptors were then blocked for 15 minutes using FcBlock (BioRad – BUF041A). Cells were pelleted by centrifugation (300 x g, 5 minutes) and resuspended in FACS buffer (100µL per 100,000 cells) and stained with fluorophore-conjugated antibodies (1:100) for 1 hour on ice. Fluorophore-conjugated antibodies (1:100) were added to whole blood for 1 hour on ice. For the final 30 minutes of staining LIVE/DEAD yellow (1:1000) (Thermo Scientific L34967) was added to cell mixtures. Cells were pelleted by centrifugation (500 x g, 5 minutes). Pellets were treated with ACK lysis buffer for 5 minutes and then resuspended in 1mL of FACS buffer (PBS, 2mM EDTA, and 1%BSA). Cells were washed in PBS (x3) after ACK Lysis with FACS buffer. Flow cytometry collection and deconvolution was performed on an Cytek Aurora Borealis 5 laser Spectral Flow Cytometer. Automatic deconvolution was performed using single stains generated from splenocytes. Gating and post-hoc analysis was performed with FCS-Express 7.18. Representative gating is presented for all flow cytometry experiments in supplemental figures.

### Antibodies

Flow cytometry antibodies used included CD45 anti-mouse PE-Cy7 (Invitrogen – 25-0451-82), CD11c rat anti-mouse Super Bright 780 (Invitrogen - 78-0114-82), CD3 anti-mouse PE (Invitrogen - 12-0032-82), CD4 anti-mouse Brilliant Ultra Violet 661 (Invitrogen – 376-0041-82), LIVE/DEAD Yellow Cell Stain for 405 (L34967), CD8a anti-mouse APC (Invitrogen – APC-65069-100UG), CD19 anti-mouse Brilliant Ultra Violet 805 (Invitrogen – 368-0193-82), CD14 anti mouse PerCP-710 (Invitrogen – 46-0141-82) and NK1.1 anti mouse FITC (Invitrogen 11-5941-82).

## Acknowledgements

Financial Support from US/DHHS/PHS/NIH: R01AI144026 (to WLS & KRL), F31AI174782 & T32NS115657 (to ARM), RF1AG079520 (to AG), R01HL148004 (to SBGA), R01DC018842 (to J-BS), R01HL137112 & R01HL165143 (to BEI), R56HL158886 (to NL), T32HL007284 & F31HL165918 (to CMP), U01AG070960 & R21OD032144 (to WX). The authors thank Bodo Levkau (Heinrich Heine University, Düsseldorf, Germany) for providing us with his laboratory’s RBC release assay protocol and Rahul Sharma (University of Virginia) for providing us with his laboratory’s splenocyte migration assay protocol.

## Author Contributions

In addition to contributing to the editing of the manuscript, individual contributions of authors:

YK experimental design, SphK assays, lymph, plasma, BAL isolation and sample preparation for LCMS, lymphocyte counting, blinded observer in EAE study, generation of multiple figures, extensive editing of manuscript, analyses of data

TH experimental design, splenocyte migration assay, *ex vivo* RBC S1P release assay, LCMS measurements and analyses of data

KD synthesis and chemical characterization of SLF80821178

DF initial synthesis and chemical characterization of SLF80821178

GMPRS surgical implantation of telemetric HR and MAP probes in rats, analyses of data

SBGA provided facilities and oversaw telemetric recording from rats, analyses of data

ARM EAE data analysis and figure generation and contributed to drafting of manuscript

AG provided facilities and oversaw EAE study, analyses of data, contributed to drafting of manuscript

CMP performed flow cytometry and subsequent data analysis and figure generation and contributed to drafting manuscript

NL provided facilities and oversaw flow cytometry analysis

KEN performed ABRs and subsequent data analysis and figure generation and contributed to drafting the manuscript

ZJJ provided cultured LECs and culture media

BEI provided facilities, bulk RNAseq data from cultured LECs and advice on LEC biology J-BS provided facilities and oversaw ABR monitoring, analyses of data

WX designed Crispr strategy and generated *Spns2*^-/-^ mouse strain

WLS provided facilities and oversaw synthetic chemistry and conception of SLF80821178 scaffold

KRL provided facilities and oversaw overall project, blinded observer in EAE study, mouse colony management, wrote manuscript

## Abbreviations

S1P: sphingosine 1-phosphate
SRM: S1P receptor modulator (agonist drug)
STB: S1P transport blocker
Spns2: Spinster homolog 2 (S1P transporter)
RBC: red blood cell
Mfsd2b: RBC & platelet S1P transporter
BAL: bronchoalveolar lavage
ABR: auditory brainstem response
LCMS: liquid chromatography mass spectrometry

